# GOntact: using chromatin contacts to infer target genes and Gene Ontology enrichments for *cis*-regulatory elements

**DOI:** 10.1101/2022.06.13.495495

**Authors:** Alexandre Laverre, Eric Tannier, Philippe Veber, Anamaria Necsulea

**Author notes:** Co-corresponding authors., **Address for correspondence,** Philippe Veber, Laboratoire de Biometrie et Biologie Evolutive, CNRS, Universite Claude Bernard Lyon 1, 43 Bd. du 11 Novembre 1918, 69622 Villeurbanne CEDEX, France, Anamaria Necsulea, Laboratoire de Biometrie et Biologie Evolutive, CNRS, Universite Claude Bernard Lyon 1, 43 Bd. du 11 Novembre 1918, 69622 Villeurbanne CEDEX, France.

## Abstract

*Cis*-regulatory elements (CREs) can be efficiently predicted genome-wide, but identifying their target genes remains challenging. Regulatory interactions between genes and CREs can take place over long genomic distances, often bypassing genes. Inferring CRE targets based on genomic proximity, as traditionally done in genomic studies, can thus be misleading. Thanks to chromosome conformation capture techniques, chromatin contacts between CREs and gene promoters can be assayed at the genome-wide scale, thus permitting more accurate predictions of CRE target genes.

Here, we present a standalone computational tool and webserver named GOntact, which infers CRE target genes using chromosome conformation capture data. GOntact can be used to derive Gene Ontology (GO) enrichments for CRE sets, thus providing a basis for functional interpretation. We apply GOntact on enhancers active in several embryonic tissues, using Promoter Capture Hi-C data to infer chromatin contacts between genes and CREs. We show that GOntact predicts functional annotations that are coherent with enhancer activity. Compared to genomic proximity, GOntact predicts consistent but more specific functional annotations.

With the increasing availability of high-resolution chromatin contact data, we believe that GOntact can provide better-informed target genes and functional predictions for CREs.

GOntact is available at https://gontact.univ-lyon1.fr (webserver) and https://gitlab.in2p3.fr/anamaria.necsulea/GOntact (standalone command-line tool).

## Introduction

In multicellular organisms, gene expression is regulated by complex interactions involving numerous *trans*-acting factors and *cis*-regulatory elements (CREs). These elements, which are by definition situated on the same chromosome as their target genes, can activate or repress gene expression (in the case of enhancers or silencers, respectively). The genomic positions and patterns of activity of enhancer elements can now be efficiently assayed with various molecular techniques, including chromatin immunoprecipitation followed by sequencing (Wei et al. 2006; Visel et al. 2009), DNase I hypersensitivity assays (Thurman et al. 2012), or assays for transposase-accessible chromatin using sequencing (Buenrostro et al. 2013). These techniques are commonly used to predict CRE positions. Genome-wide maps of CREs are now readily available, in particular for model organisms (ENCODE Project Consortium et al. 2020). However, identifying CRE target genes remains a difficult question, even when their localization and molecular function have been ascertained.

The most commonly used approach to predict CRE target genes is based on an assumption of genomic proximity (McLean et al. 2010). This assumption is undoubtedly reasonable, at least for enhancer elements, which were originally demonstrated to act on their neighboring genes (Banerji et al. 1983). However, there is now solid evidence that enhancers can be situated at large genomic distances from the genes they regulate. A now classical example is the ZRS enhancer of *Shh*, which is situated 1 megabase (Mb) away from its target gene (Lettice et al. 2002). Through fluorescence *in situ* experiments, it was demonstrated early on that distal enhancers are brought in physical proximity with target gene promoters in the nucleus, likely through the formation of chromatin loops (Carter et al. 2002; Amano et al. 2009). The physical proximity between DNA segments that are distant on the linear genome can be detected through chromosome conformation capture techniques (Dekker et al. 2002; Lieberman-Aiden et al. 2009). Applications of these techniques have confirmed that long-range regulatory interactions between enhancers and target genes are frequent (Montavon et al. 2011; Smemo et al. 2014; Schoenfelder et al. 2015). Additional evidence comes from perturbation experiments, which indicated that a large fraction of regulatory interactions (up to 30%) occur between enhancers and genes that are not their nearest neighbors (Gasperini et al. 2019).

Recent technical developments have enabled high-resolution mapping of chromatin contacts between promoters and distal CREs. The Hi-C approach was designed to investigate chromatin contacts between all possible pairs of genomic regions, at a genome-wide scale (Lieberman-Aiden et al. 2009). However, this approach requires great sequencing depth to achieve high resolution and sensitivity (Parker et al. 2023). The Micro-C technique, in which the genome is digested with a micrococcal nuclease rather than with restriction enzymes (Hsieh et al. 2015), enables better resolution (Lee et al. 2022). Other Hi-C variants offer better resolution and sensitivity for a restricted class of chromatin interactions, such as those mediated by a given protein or in which the interacting regions carry a specific histone mark (Mumbach et al. 2016). For promoter-CRE interactions, capture Hi-C and capture Micro-C approaches are particularly well suited, as they can be designed to target chromatin contacts that involve gene promoters, without additional requirements on the chromatin modifications of the promoter or on protein mediators (Mifsud et al. 2015; Schoenfelder et al. 2015; Golov et al. 2020; Downes et al. 2021; Lee et al. 2022). The resulting data is enriched in regulatory interactions between promoters and CREs and can thus provide a better-informed prediction of CRE target genes.

To take advantage of the increasing availability of high-resolution chromatin contact data, we developed a new tool named GOntact, which uses this type of data to infer associations between CREs and genes. Importantly, GOntact can be used to derive gene ontology (GO) enrichments for sets of CREs from these associations, thus providing a basis for functional interpretation. Here, we apply GOntact on experimentally validated enhancer datasets for human and mouse, using Promoter Capture Hi-C (PCHi-C) data for chromatin contact inference. We show that GOntact generates functional annotations that are coherent with the activity patterns of the enhancers. We find that there is substantial overlap between functional predictions obtained based on genomic proximity (McLean et al. 2010) and based on chromatin contacts, but that the latter approach pinpoints more specific GO categories. GOntact can thus prioritize testable hypotheses for CRE functional characterization.

## Results

### Predicting CRE targets and functions based on chromatin contacts with GOntact

We developed GOntact, a tool that aims to provide an easy means of predicting CRE target genes based on chromatin contacts. Given a set of CREs, a genome annotation and a list of chromatin contacts, GOntact identifies chromatin contacts that link gene promoters to CREs (Materials and Methods, Supplementary Figure 1). The GOntact tool has two sub-commands: the first one (“gontact annotate”) outputs a list of CRE-gene associations, including the evidence on which the association was inferred; the second one (“gontact enrich”) performs a gene ontology (GO) enrichment analysis for a given set of CREs, with respect to a background set of CREs or to the entire genome (Materials and Methods). GOntact enables filtering of the input chromatin contact dataset and allows users to fine-tune the parameters used for gene-CRE associations (Materials and Methods).

To enable comparisons between results obtained with chromatin contacts or based on genomic proximity, we re-implemented within GOntact the main principle of the widely used GREAT (Genomic Regions Enrichment of Annotations Tool) method (McLean et al. 2010). GOntact also enables using a simple genomic proximity approach, in which CREs are associated to genes if the distance to gene promoters is below a user-defined threshold (hereafter, this method is referred to as the “fixed window” approach). Users can switch between the approaches based on chromatin contacts or based on genomic proximity by setting the “mode” argument in the command line.

### Inferring CRE target genes based on chromatin contacts and genomic proximity

We illustrate the different approaches for predicting CRE target genes with GOntact focusing on the genomic region around the human *POU6F1* gene (Figure 1A), for ENCODE-predicted enhancers. The simplest inference method based on genomic proximity attributes to a given gene all CREs found in its vicinity, that is, within a fixed window centered on the gene transcription start site (TSS). In the widely used GREAT (Genomic Regions Enrichment of Annotations Tool) method, a more complex principle is proposed (Figure 1A). For each gene, a basal regulatory domain is constructed, which spans a certain distance upstream and downstream of the gene TSS (default values 5 kilobases (kb) and 1 kb, respectively). These domains are then extended on each side, at most reaching the basal regulatory domains of the neighboring genes (on either DNA strand), and no further than a given genomic distance (1 Mb by default). As illustrated in Figure 1A, this results in regulatory domains of variable sizes, depending on the local gene density. The associations between genes and CREs depend on the quality of the genome annotations: a gene missing from or wrongly included in the genomic annotation may affect the attribution of CREs to its neighbors. Of note, with this approach, genes are previously filtered to discard those that lack Gene Ontology annotations.

**Figure 1.**
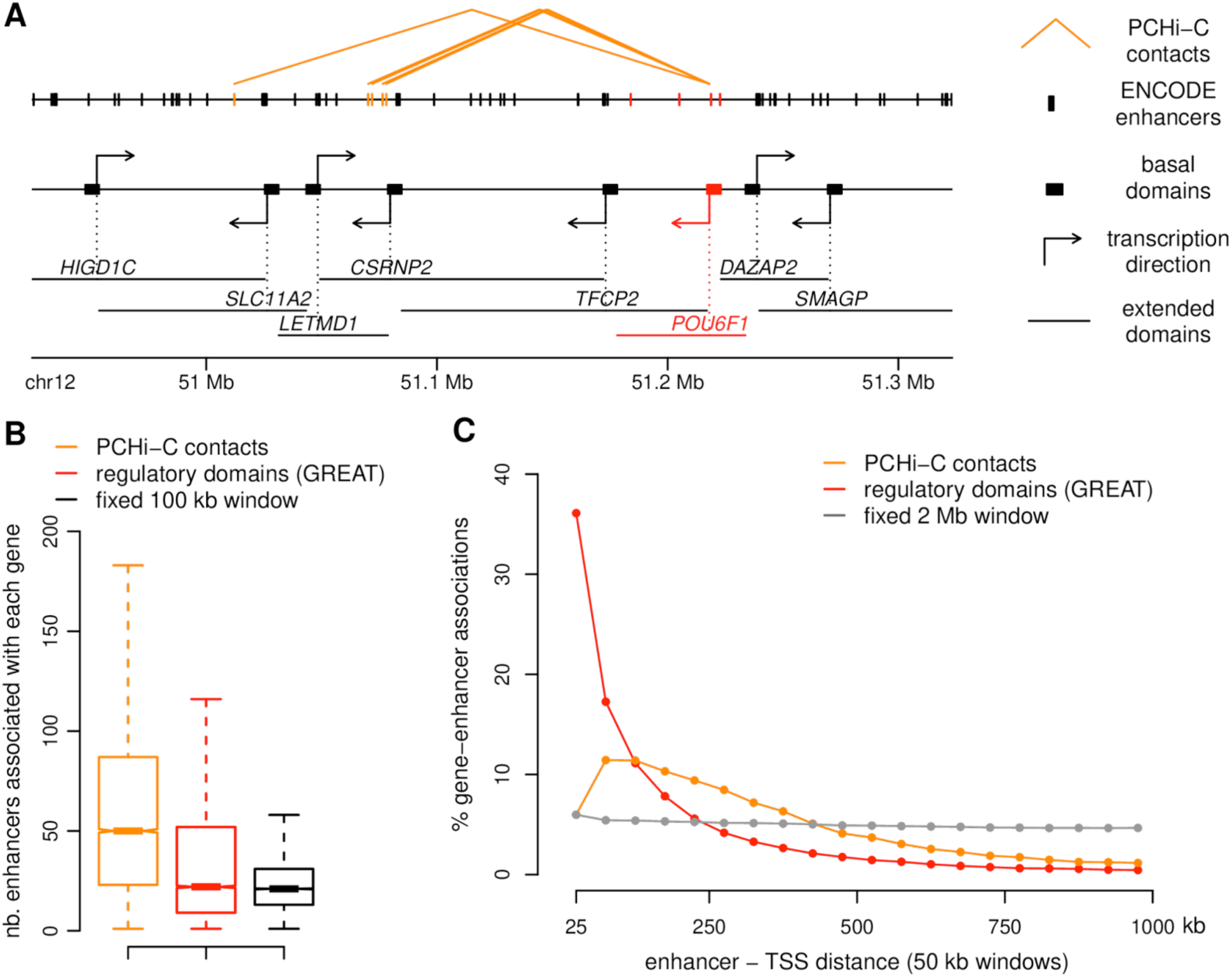
Gene-enhancer associations. **A)** Illustration of gene-enhancer associations around the *POU6F1* locus in the human genome. Top panel: position of predicted enhancers (ENCODE) and depiction of PCHi-C chromatin contacts between the *POU6F1* gene promoter and predicted enhancers. Orange rectangles represent enhancers associated with the *POU6F1* gene through PCHi-C contacts, red rectangles represent enhancers associated with the *POU6F1* gene through the “regulatory domains” (GREAT) approach (McLean et al., 2010). Second panel: depiction of the basal regulatory domains proposed by McLean et al., 2010, comprising 5 kb upstream and 1 kb downstream of the transcription start site. Third panel: depiction of extended regulatory domains, which are limited by the basal regulatory domains of upstream and downstream neighboring genes, within a maximum distance of 1 Mb. Only protein-coding genes are shown. **B)** Boxplots representing the distribution of the number of enhancers associated with each gene, through PCHi-C contacts (orange), regulatory domains (red) and simple genomic proximity (fixed 100 kb window, black). Only genes associated with enhancers with all three approaches were considered. **C)** Distribution of the distances between genes and associated enhancers, as predicted with different approaches (PCHi-C contacts, regulatory domains and fixed 2 Mb window). For this panel, the size of the fixed window was set to 1 Mb for consistency with GOntact and GREAT.

In contrast, CRE target gene predictions based on chromatin contacts do not depend on the quality of genome annotations, but instead likely rely on the quality of the chromosome conformation capture data used as input. In Figure 1A, we illustrate the enhancer elements that are attributed to *POU6F1* by GOntact, based on PCHi-C data derived from multiple cell types and conditions (Laverré et al. 2022). They are situated more than 100 kb away from the *POU6F1* TSS and bypass at least two other gene promoters. With the genomic proximity approach (GREAT regulatory domains), 4 enhancers are associated with *POU6F1,* at distances ranging from 700 bp to 34 kb (Figure 1A). There is no overlap between the sets of CREs attributed to *POU6F1* with the two approaches. This may be in part due to the fact that the PCHi-C approach cannot efficiently detect short-range contacts.

Overall, individual genes are assigned substantially more enhancers based on PCHi-C contacts than based on genomic proximity (Figure 1B). The median number of ENCODE-predicted enhancers assigned to a gene is 22 with GREAT and 50 with GOntact, for human (Wilcoxon rank sum, p-value *<* 1e-10, Figure 1B). The median number of enhancers attributed to genes is similar between GREAT and a simple fixed 100 kb window approach (21 enhancers), but the two distributions are significantly different (Wilcoxon rank sum, p-value *<* 1e-10, Figure 1B). Conversely, the distributions of the numbers of genes associated with each enhancer are also significantly different between the three approaches (Wilcoxon rank sum, p-value *<* 1e-10 for all three comparisons, Supplementary Figure 2A). With GOntact, 55% of enhancers are associated with at most 2 genes, while with GREAT this is true for 99% of enhancers, by construction (extended regulatory domains cannot overlap among more than two genes). Importantly, the sets of enhancers attributed to each gene differ considerably among methods. The median Jaccard index (which measures the ratio between the size of the intersection and the size of the union of two sets) is equal to 0.01 for the comparison between GREAT and GOntact (Supplementary Figure 2B). There is more similarity between GREAT and the fixed 100 kb window approach (median Jaccard index 0.35, Supplementary Figure 2B), as expected given that both methods are based on genomic proximity.

The genomic distances between promoters and enhancers predicted to be in regulatory interactions vary considerably among methods. With GREAT, 53% of all gene-enhancer pairs are separated by less than 100 kb and only 8% are separated by more than 500 kb (Figure 1C). With GOntact, distances between pairs of enhancers and promoters are generally larger: only 18% are separated by less than 100 kb, and 20% are separated by more than 500 kb (Figure 1C). In contrast, regulatory associations inferred from PCHi-C data are less frequent in close proximity to genes, which may be due to the technical difficulty of distinguishing genuine from spurious chromatin contacts at small distances on the linear genome (Cairns et al. 2016).

### Gene Ontology enrichment analyses for *cis*-regulatory elements

Gene Ontology (GO) enrichment analyses contrast a focal gene set (*e.g.* genes that are up-regulated in a given condition) with a background gene set (*e.g.* all analyzed genes). They are traditionally used as a basis for functional predictions. Such analyses can be used to formulate new hypotheses that can be validated through further experiments. Similar analyses can be performed for sets of CREs, by transferring GO annotations to CREs from the genes they are predicted to regulate. This principle has been implemented within GREAT (McLean et al. 2010). We applied the same principle in GOntact, by transferring GO categories to CREs from the genes with which they form chromatin contacts (Materials and Methods).

We used three different methods to infer gene-CRE associations and thereby GO-CRE associations: GOntact applied on PCHi-C chromatin contacts, GREAT regulatory domains, and a simple genomic proximity method (CREs are attributed to genes if found within a fixed 100 kb window around the TSS). We performed GO enrichment analyses for three sets of experimentally validated enhancers, which are active in the embryonic forebrain, limb or heart. Enhancer coordinates and activity patterns in embryonic tissues were downloaded from the VISTA Enhancer browser (Kosicki et al. 2025). We performed GO enrichment analyses contrasting each of these enhancer sets with the full set of ENCODE-predicted enhancers (Materials and Methods). This comprehensive enhancer dataset was obtained using a large number of cell types, tissues and biological conditions. Alternatively, we performed GO enrichment analyses using the whole genome as a background, with similar results (Supplementary Dataset 1; Materials and Methods).

Overall, we found good agreement among the three methods for the top GO categories (Figure 2, Supplementary Figures 3-4). The top GO categories were related to neuron differentiation and brain development for forebrain enhancers (Figure 2), to digit and limb morphogenesis for limb enhancers (Supplementary Figure 3), and to cardiac muscle development for heart enhancers (Supplementary Figure 4). All three methods thus propose functional categories that are consistent with enhancer activity patterns. However, we note that the top GO categories put forward by each method are not identical. For example, for forebrain enhancers, the 2nd GO category found significant by GOntact is “forebrain radial glial cell differentiation”, which does not appear in the top 15 GO categories with the two other approaches (Figure 2). Conversely, the two approaches that rely on genomic proximity show significant enrichment of GO categories related to transcriptional regulation, which do not appear at the top of the list for GOntact (Figure 2). Likewise, for embryonic heart enhancers, GOntact tends to bring forward categories related to the early development of the organ (e.g., lateral mesoderm formation, embryonic heart tube elongation), whereas genomic proximity methods seems to prioritize categories related to the physiology of the embryonic and adult organ, such as the regulation of striated muscle contraction or the regulation of blood circulation (Supplementary Figure 4).

**Figure 2.**
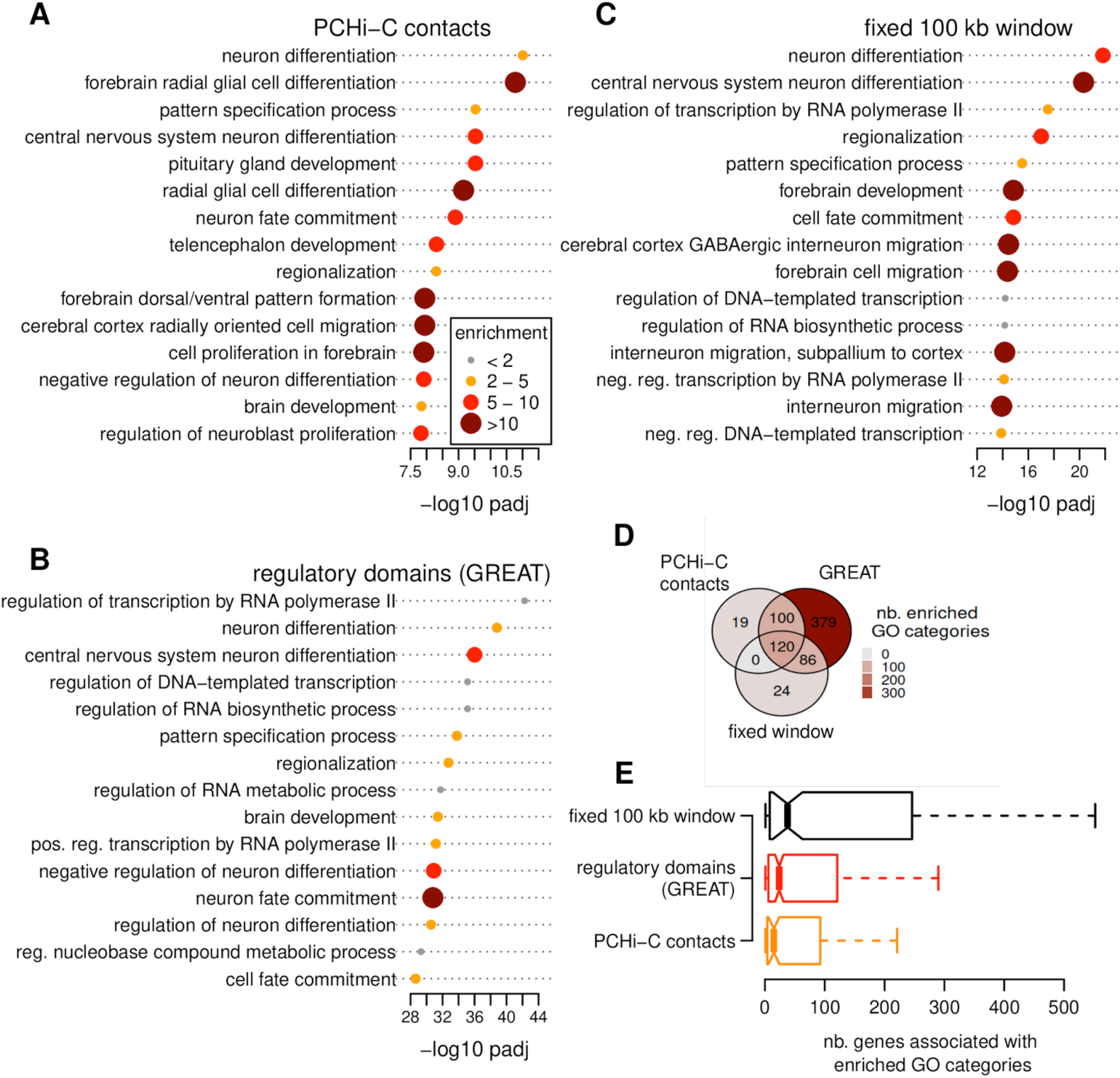
Gene Ontology enrichments for Vista forebrain enhancers. **A)** Barplot representing log10-transformed adjusted p-values for the top 15 significantly enriched GO categories, for a GO enrichment analysis performed based strictly on PCHi-C chromatin contacts (we did not include a basal regulatory domain). Dot sizes and colors represent enrichment values. **B)** Same as **A)**, for the “regulatory domains” approach (McLean et al., 2010). **C)** Same as **A)**, for the simple genomic proximity approach (fixed 100 kb window). **D)** Venn diagram representing the overlap between the GO categories found to be significantly enriched (FDR < 0.01) with the three approaches. **E)** Boxplot representing the number of genes associated with enriched GO categories, for the three methods. For all panels, enrichment analyses are performed using as a background the set of human ENCODE-predicted enhancers. Analyses performed using the whole genome as a background are provided in Supplementary Dataset 2.

The numbers of significantly enriched GO categories (FDR < 0.01) differ considerably among the three methods, with far larger numbers observed with GREAT for all three enhancer sets (Figure 2, Supplementary Figures 3-4). Interestingly, the enriched GO categories are generally associated with larger numbers of genes for the two methods based on genomic proximity than for GOntact (Figure 2, Supplementary Figures 3-4). This suggests that GOntact can prioritize more specific biological processes.

Consistent with the observations on the top 15 GO categories, we observed that overall, the ranks of the enriched GO categories (ordered based on the adjusted p-value associated with the GO enrichment test) are only moderately correlated among the three approaches (Supplementary Figure 5). However, the enrichment values (*i.e.,* the ratio between the frequencies of the GO categories in the foreground and background CRE sets) are well correlated among methods, with Spearman’s rho coefficients ranging from 0.9 to 0.97 (Supplementary Figure 6).

### Influence of GOntact parameters on GO enrichment results

To test how GOntact parameters affect GO enrichment results, we extracted two relevant GO categories for each tissue (*i.e.* those that were at the top of the enrichment lists with default parameters: “neuron differentiation” and “regulation of transcription by RNA polymerase II” for forebrain enhancers, “limb morphogenesis” and “embryonic digit morphogenesis” for limb enhancers, “heart development” and “muscle tissue morphogenesis” for heart enhancers). We then analyzed their enrichment values and significance levels for GOntact runs with different parameters (Figure 3, Supplementary Figure 7).

**Figure 3.**
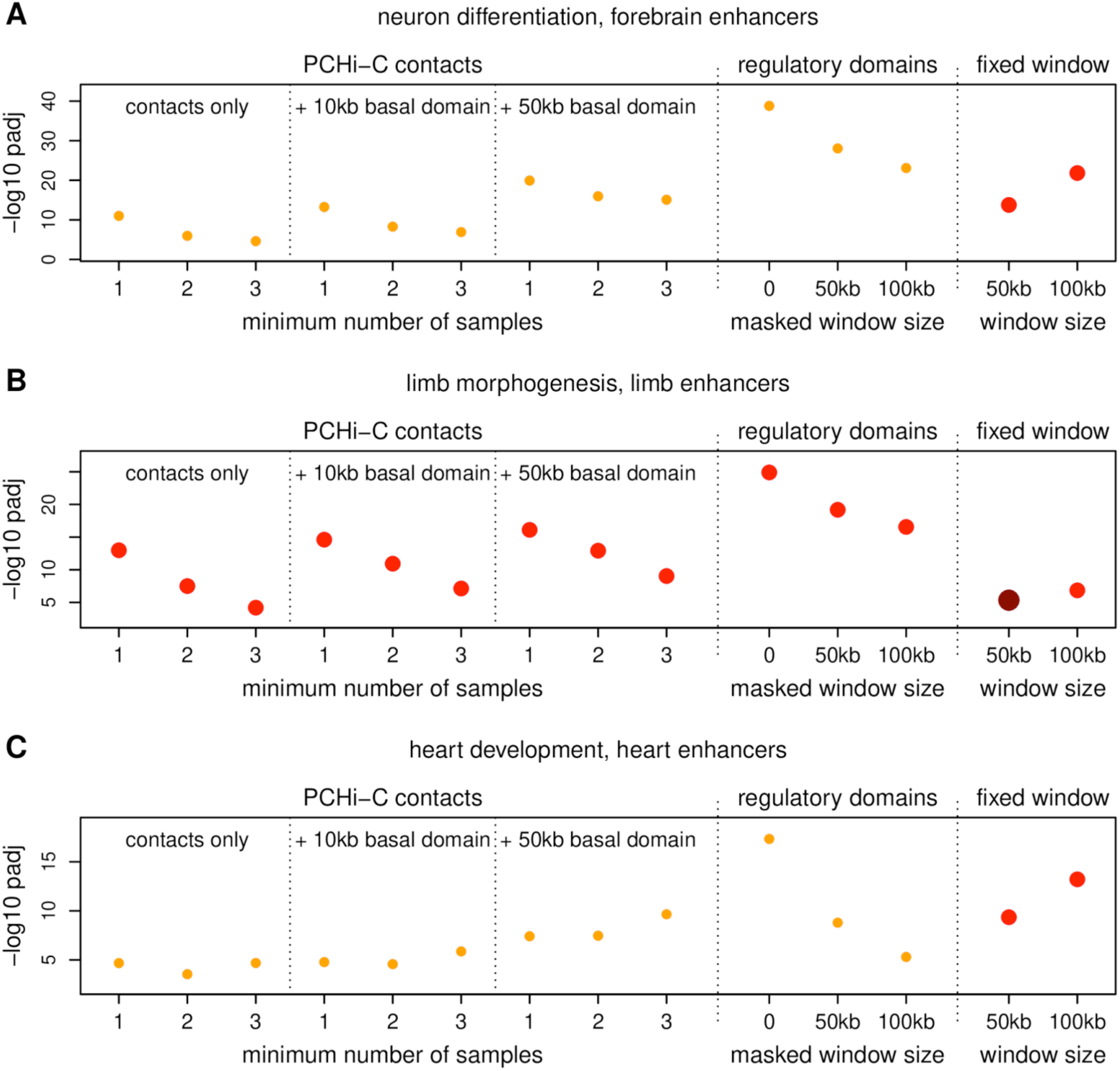
GO enrichment results obtained with different approaches and parameter sets. **A)** Dot plot representing the log10-transformed adjusted p-value for the “neuron differentiation” GO category, for GO enrichment analyses contrasting human forebrain enhancers (Vista) with ENCODE-predicted enhancers. Different columns represent method variants, in order: PCHi-C contacts, requiring that chromatin contacts be observed in at least 1, 2 and 3 samples, respectively; PCHi-C contacts + basal 10 kb domain, with contacts observed in at least 1,2 and 3 samples; PCHi-C contacts + basal 50 kb domain, with contacts observed in at least 1,2 and 3 samples; regulatory domains (basal + extended domains, maximum 1 Mb); modified regulatory domains, in which a 50 kb window centered on the TSS is “masked”; modified regulatory domains, in which a 100 kb window centered on the TSS is “masked”; fixed 50 kb window centered on the TSS; fixed 100 kb window centered on the TSS. **B)** Same as **A)**, for the “limb morphogenesis” GO category, for human limb enhancers. **C)** Same as **A)**, for the “heart development” GO category, for human heart enhancers. Dot sizes and colors represent enrichment values, as in Figure 2.

We first wanted to evaluate how the quantity and the quality of the input chromosome conformation capture data may affect the results. The GOntact results presented above were performed using chromatin contacts derived from 26 human PCHi-C samples (Laverré et al. 2022). These included numerous sample-specific contacts. We reasoned that contacts shared across multiple biological samples may be more relevant to define target genes for sets of CREs that are derived from unrelated tissues, as these shared contacts may include pre-formed interactions observed in the absence of CRE and gene activity (Paliou et al. 2019). Overall, similar GO enrichment results were observed when requiring contacts to be observed in 2 or 3 samples (Supplementary Dataset 1). However, we observed a decrease in detection sensitivity: the significance level associated with the most relevant GO categories decreased in this setting (Figure 3, Supplementary Figure 7). The decrease in sensitivity is likely due to the fact that fewer chromatin contacts are used for target gene prediction. Interestingly, this pattern was not observed for all GO categories: for the “heart development” GO category, a more significant enrichment was observed with shared PCHi-C contacts (Figure 3).

Because significant interactions at short genomic distances are depleted in chromatin contact data, the default parameters of GOntact exclude chromatin contacts occurring at distances below 25 kb. As there is a genuine enrichment for CREs in close proximity to gene promoters (Gasperini et al. 2019), we wanted to test whether adding a proximal regulatory domain to PCHi-C contacts could improve our results (Materials and methods). Overall, the significance levels of the GO enrichment tests for the relevant GO categories were stronger upon adding a 10 kb or a 50 kb basal domain centered around the TSS, for the three analyzed tissues (Figure 3, Supplementary Figure 7).

We next wanted to test to what extent the GO enrichment results obtained with GREAT can be explained by the abundance of CREs in the immediate vicinity of gene TSS. To do this, we masked a genomic region centered on the TSS in the GREAT analysis. That is, rather than incorporating a basal regulatory domain, we exclude gene-CRE associations at distances smaller than a certain threshold (25 kb or 50 kb in this case, Figure 3, Supplementary Figure 7). We observed that the significance of the GO enrichment test was lower following the masking procedure, for all analyzed GO categories and for all three enhancer sets. Notably, the drop was larger for some GO categories, such as “heart development” for embryonic heart enhancers. Conversely, the fixed window approach generally provided stronger significance values for larger window sizes, although we note that the increase in statistical significance was only moderate for some GO categories (e.g., “limb morphogenesis” for embryonic limb enhancers, Figure 3).

### GO enrichment for human-specific conserved element deletions (hCONDELs)

Finally, we applied GOntact to a set of ∼10,000 conserved elements that experienced human-specific deletions (Xue et al. 2023). These elements were named hCONDELs (Xue et al. 2023). Sequences that are conserved across large evolutionary distances but accelerate specifically in the human lineage have attracted considerable attention, as they may include elements that are responsible for the emergence of human-specific phenotypes (Pollard et al. 2006; McLean et al. 2011). Many of these elements are involved in gene expression regulation (Capra et al. 2013). An analysis performed with GREAT showed that genes near hCONDELs are enriched in transcriptional regulatory functions, but also in neural development (Xue et al. 2023). We performed GO enrichment analyses on this set of elements with the three approaches described above, using the whole genome as a background. As observed for VISTA enhancer analyses, overall the three methods were in good agreement, confirming the enrichment in GO categories associated with transcriptional regulation and with developmental processes (Figure 4). Here again, the three approaches highlight different GO categories at the top of the enrichment results. Notably, among the top 15 GO categories obtained with GOntact, two (ranked 1st and 9th) are associated with cranial nerve morphogenesis. The top GO category is related to the morphogenesis of the glossopharyngeal nerve, which has sensory and motor functions associated with the pharynx and the larynx. This specific GO category may be relevant for the emergence of vocalization-related processes. In contrast, the GO categories that are at the top of the significance list with approaches based on genomic proximity are more general, and relate to the regulation of transcription or to system development. Thus, in this situation, GOntact can provide a different list of candidates for further functional exploration than proximity-based approaches.

**Figure 4.**
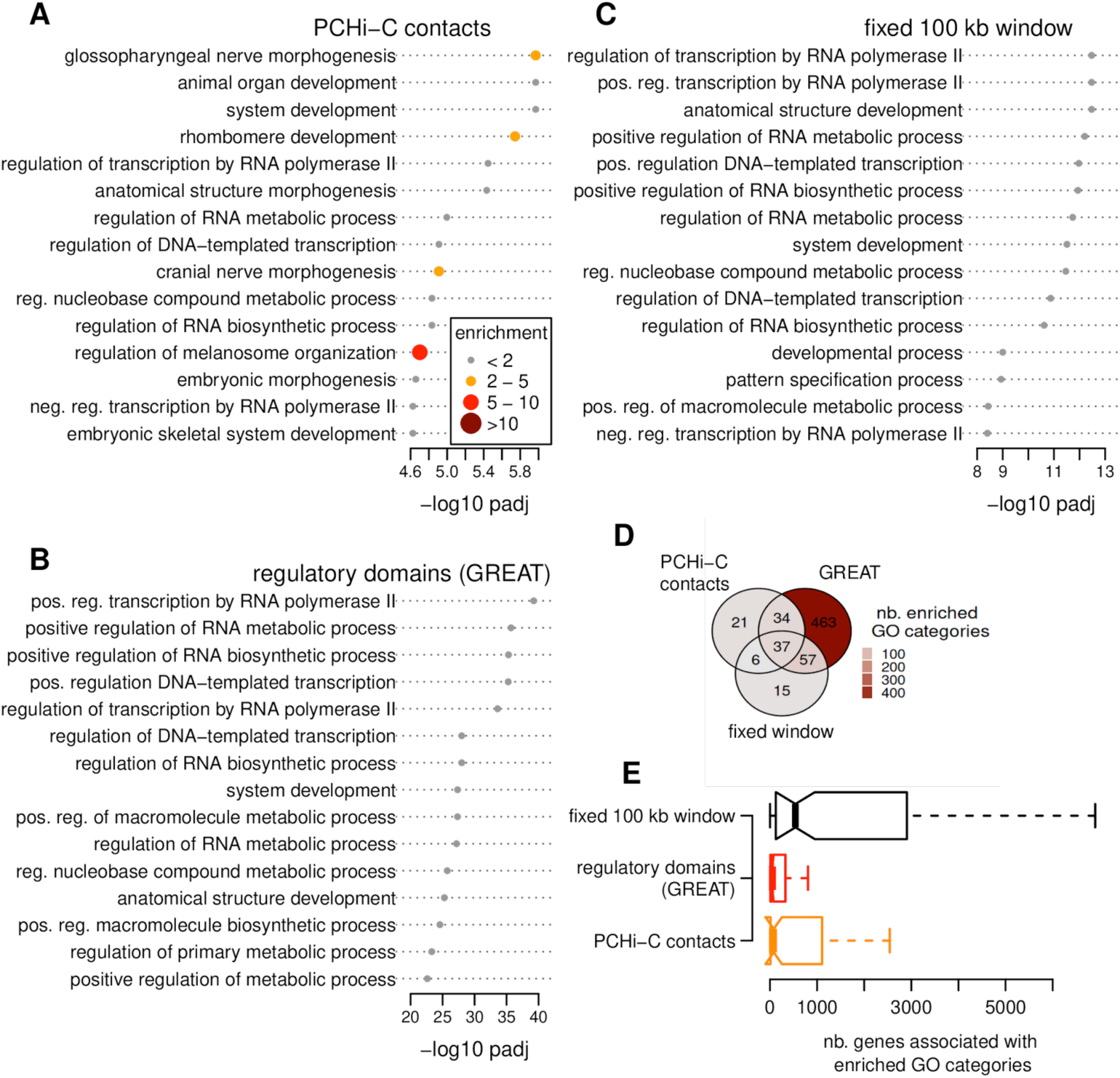
Gene Ontology enrichments for human-specific conserved element deletions (hCONDELs). **A)** Barplot representing log10-transformed adjusted p-values for the top 15 significantly enriched GO categories, for a GO enrichment analysis performed based strictly on chromatin contacts (we did not include a basal regulatory domain). Dot sizes and colors represent enrichment values. **B)** Same as **A)**, for the “regulatory domains” approach (McLean et al., 2010). **C)** Same as **A)**, for the simple genomic proximity approach (fixed 100 kb window). **D)** Venn diagram representing the overlap between the GO categories found to be significantly enriched (FDR 0.01) by the three approaches. **E)** Boxplot representing the number of genes associated with enriched GO categories, for the three methods. For all panels, enrichment analyses are performed using the whole genome as a background. **A-C)** Dot sizes and colors represent enrichment values.

## Discussion

Here, we propose a new tool for CRE target gene predictions and functional enrichment analysis, which uses chromatin contacts as a basis. Evaluating the performance of GOntact in a purely computational setting is a difficult task, as no ground truth is available. Nevertheless, we rely on an extensive body of literature, which demonstrates that promoter-CRE chromatin contacts are enriched in genuine regulatory interactions (Schoenfelder et al. 2015; Mifsud et al. 2015; Javierre et al. 2016; Jung et al. 2019). We performed all analyses with GOntact and GREAT, a widely used software that implements CRE target gene predictions based on genomic proximity.

Both GOntact and GREAT aim to predict putative biological functions for a set of *cis*-regulatory elements of gene expression. We applied these methods on sets of experimentally validated enhancer elements, whose pattern of activity was ascertained in transgenic mouse embryos (Kosicki et al. 2025). We thus expect an enrichment for biological processes associated with the development and the physiology of the organs in which enhancer activity was detected. Reassuringly, all the methods that we tested here show results that are coherent with these expectations. GOntact and GREAT provide consistent Gene Ontology enrichment results, although these two methods rely on very different approaches for CRE target gene inference. We believe that this is due to the fact that both methods capture functionally relevant gene-CRE associations: GREAT perfectly captures short-distance regulatory relationships, which represent the majority of gene-CRE associations (Gasperini et al. 2019), while GOntact captures long-distance regulatory interactions. We thus recommend that both types of methods be applied and compared, as they each may prove to be relevant in different contexts.

Beyond the overall agreement between GOntact and proximity-based approaches, some differences are notable. In particular, we find that proximity-based approaches output larger numbers of significantly enriched GO categories, but that these categories are more general (that is, they are associated with larger numbers of genes) than those put forward by GOntact. This is particularly evident in the hCONDELs analysis, where the GO categories at the top of the enriched list pinpoint specific functions related to cranial nerve morphogenesis, while proximity-based approaches highlight more general functions related to transcriptional regulation. The fact that GOntact outputs narrower lists of enriched GO categories may be an advantage in exploratory analyses, as they can help in the decision process for further experimental validation.

A further advantage of the regulatory relationships inferred by GOntact is that they are to a large extent independent of the organization of the genome and of the quality of our knowledge thereof, in contrast with the methods implemented in GREAT. In well-annotated genomes such as human and mouse, GREAT undoubtedly performs very well. However, genome annotation quality can affect the target gene inference as performed by GREAT. A missing gene or a wrongly assigned transcription start site can have an inordinate impact on the definition of the regulatory domains of neighboring genes, and thus on CRE target gene prediction.

While GREAT is dependent on genomic organization and on the quality of genomic annotations, GOntact is evidently dependent on the quantity, quality and specificity of the input data, in particular the set of PCHi-C chromatin contacts. Here, we used a previously published set of chromatin interactions, derived from multiple cell types and conditions (Laverré et al. 2022). A non-negligible fraction of these interactions were only observed in a single cell type (Laverre et al. 2022). Ideally, chromatin interactions should be provided for the same biological conditions (tissues, cell types, developmental stages etc.) in which CRE activity is assessed. In the absence of perfectly matched chromatin contact and enhancer activity data, one solution is to use as an input a set of chromatin interactions that are shared across multiple biological conditions, which are thus more likely to be relevant in the biological context that is being examined. However, we note that this filter greatly reduces the number of usable interactions and thus results in a decrease in sensitivity.

Finally, we note that generating chromatin contact data is becoming increasingly accessible. Hi-C experiments are now typically performed for genome assembly projects and for analyses of broad-scale chromosome conformation features, including topologically associating domains (Nora et al. 2012). While these data lack the resolution and sensitivity of PCHi-C and of other capture-based approaches, with sufficient sequencing depth they provide chromatin contacts at the kilobase scale (Lee et al. 2022). Several computational methods were proposed to detect chromatin loops from Hi-C data (Kaul et al. 2020; Lu et al. 2021; Lagler et al. 2021), which can be used as an input for GOntact. We thus believe that GOntact can provide a practical alternative to approaches based purely on genomic proximity for CRE target enrichment and functional enrichment analyses.

## Materials and methods

### The GOntact tool and webserver

GOntact is a standalone command-line tool developed in OCaml. It can be installed using the OCaml package manager (opam), or by directly compiling the source code. Alternatively, it can be used through Docker and Apptainer containers. The source code, the Docker container, as well as detailed installation and usage instructions, are available in a public GitLab repository:

https://gitlab.in2p3.fr/anamaria.necsulea/GOntact

Manual pages are accessible with the “--help” argument for the command line tool.

A GOntact webserver is also available (https://gontact.univ-lyon1.fr). The webserver enables quick GO enrichment analyses for human and mouse, using chromatin contact data, genome annotations and gene ontology data stored on the server. While the server is limited at the moment to these two species, the command line tool can be used for any species for which all required data types are available. A schematic description of the required input files and of the output generated by GOntact is given in Supplementary Figure 1.

### Chromatin contact data processing

Chromatin contact data can be provided to GOntact in iBed and/or washU formats (defined in the CHiCAGO vignette (Cairns et al. 2016) and further explained in the GOntact GitLab repository). This can be done for multiple samples at the same time, by specifying several input files separated by a comma (“ibed-path” and “washu-path” arguments). GOntact enables filtering chromatin contacts based on the minimum and maximum distance between interacting regions (“min-dist-contacts” and “max-dist-contacts” arguments, default values 25 kb and 1 Mb, respectively), on the minimum interaction score (“min-score” argument, default value 5.0, which corresponds to the score cutoff proposed in ChICAGO (Cairns et al. 2016)) and on the minimum number of samples in which the contact is observed (“min-samples” argument, default value 1).

For the analyses presented in this manuscript, we ran GOntact using PCHi-C data from a previous study, which processed and combined 26 human and 14 mouse samples (Laverré et al. 2022). The PCHi-C technique aims to identify chromatin contacts between a predefined set of restriction fragments, that are “baited” using RNA molecules, and the entire genome. Baited restriction fragments were selected to target gene promoters (Mifsud et al. 2015; Schoenfelder et al. 2015). Interactions were scored using the HiCup and ChICAGO tools, on the human hg38 and mouse mm10 genome assemblies (Wingett et al. 2015; Cairns et al. 2016).

### Chromatin contact annotation

Genome annotations have to be provided to GOntact as GTF (General Transfer Format) files. They are used to annotate the gene promoters that are involved in chromatin contacts. For PCHi-C data, gene promoters are targeted by the “baited” restriction fragments, whose coordinates are provided in the first three columns of the iBed and WashU files. Following this principle, GOntact considers that the first genomic interval provided for each chromatin contact is the “baited” region. It then annotates chromatin contacts by identifying the transcription start sites that are found close to the coordinates of the first genomic interval, within a maximum distance defined by the user (default value 1 kb). For chromatin contact data that does not have “baited” regions, GOntact can force chromatin contacts to be symmetrical. Specifically, if a contact is listed between genomic regions A and B, with the “force-symmetry” option GOntact will add a contact between genomic regions B and A, if this contact is not already present in the list. Conversely, GOntact can discard interactions that occur between two “baited” genomic regions (default; the remove-bait-bait option can be set to “false” to disable this behavior).

For gene ontology (GO) enrichment analyses, only genes and transcripts annotated as “protein-coding” are retained, as GO annotations are still generally lacking for non-coding genes. For CRE target gene predictions, this behavior can be controlled with the “filter-biotypes” argument (possible values “protein_coding” or “all”, default value is “protein_coding”).

In this manuscript, we used genome annotations provided by the Ensembl database release for human hg38 (Ensembl release 115) and mouse mm10 (Ensembl release 102) genome assemblies.

### Associating *cis*-regulatory elements and target genes

The first function of GOntact is to associate *cis*-regulatory elements (CREs) with potential target genes. This function is accessible through the “gontact annotate” sub-command.

In the default run mode (“contacts”), GOntact associates CREs to genes based on chromatin contacts. CRE coordinates have to be provided to GOntact in BED format. GOntact identifies CREs that are close to genomic intervals involved in chromatin contacts, within a certain maximum genomic distance defined by the user. Only the second genomic interval provided for each chromatin contact is used, following the principle described above for PCHi-C data. For chromatin contact data that does not have “baited” regions, GOntact can enforce chromatin contacts to be symmetrical. In addition to chromatin contacts, GOntact will associate CRE to genes if the CREs are found in a basal regulatory domain around the gene TSS, consisting of an upstream interval and a downstream interval whose size is defined by the user (“upstream” and “downstream” arguments, default values 5 kb and 1 kb respectively).

GOntact also provides a “proximity” run mode, in which the association between CREs and genes is performed based on genomic proximity. In this mode, first a basal regulatory domain is defined, as described above and as proposed by McLean et al. 2010. The basal regulatory domain is then extended on both sides of the TSS, for a maximum distance defined by the user (“extend” argument, default value 1 Mb) and at most up to the boundary of the basal regulatory domain of the neighboring genes. This corresponds to the “basal + extension” run mode of GREAT (McLean et al. 2010). To associate CRE to genes with a simple genomic proximity approach, in which CREs are attributed to all genes whose TSS is found within a given maximum genomic distance, the “upstream” and “downstream” argument should be set to the desired maximum distance, and the “extend” argument should be set to 0.

To test the influence of the immediate neighborhood of the gene TSS on CRE functional annotation, we enabled the possibility of masking a genomic interval centered on the gene TSS in “proximity” mode, for GO enrichment analyses. The size of the masked genomic region can be defined with the “masked-size” argument (default value is 0, corresponding to no masking).

For regulatory domain definition, only one TSS is used for each protein-coding gene. This TSS corresponds to the isoform with the longest exonic length (including coding and untranslated regions).

### Gene Ontology enrichment analyses

The second function of GOntact is to perform gene ontology (GO) enrichment analyses for CREs. This function is accessible through the “gontact enrich” sub-command.

To use this function, GOntact users must provide a file containing the structure of the GO database (“ontology” argument), in OBO format. They must also provide a file containing the GO annotation of the studied species (“functional-annot” argument), in GAF format.

In this manuscript, we downloaded the GO database and GO annotations for human and mouse from geneontology.org, for the 2025-10-10 database release. For human we used annotations generated by the EBI Gene Ontology Annotation database and for mouse we used annotations generated by Mouse Genome Informatics, also from the 2025-10-10 release of the GeneOntology database. We used the “basic” version of the ontology, which is acyclical.

GO annotations are typically provided only for the most precise term of a given ontology branch, but genes are assumed to inherit the annotations of all the parent terms. GOntact processes the structure of the ontology and the provided gene annotations to propagate annotations through the term hierarchy. The GO domain to be analyzed can be specified with the “domain” argument (possible values “biological_process”, “molecular_function”, “cellular_component”; default value “biological_process”). In the analyses presented in the manuscript, we focused on the “biological process” Gene Ontology domain.

GO annotations are transferred to CREs from their predicted target genes. Gene-CRE associations can be obtained based on chromatin contacts or on genomic proximity, as described above. A given CRE can be associated only once with a given GO category, even if it is associated with several genes that have this GO annotation.

GO enrichment analyses are performed by comparing a foreground set of CREs (“foreground” argument, compulsory) with a background set of CREs (“background” argument, optional). If the background set is provided, the expected frequencies of GO categories are the frequencies observed in the background set.

If the background set is not provided, the expected frequencies of GO categories are computed on the entire set of chromatin contacts or on the entire set of regulatory domains, depending on the selected mode. GOntact will compute for each GO category the total number of nucleotides that are associated with this GO category. In “contacts” mode, we consider that individual nucleotides are associated with a GO category if they belong to genomic intervals that are in contact with genes associated with this category, or to basal regulatory domains associated with these genes (if a basal regulatory domain was defined with the “upstream” and “downstream” arguments). The background GO frequency is then computed as the ratio between the number of nucleotides associated with the focal GO category and the number of nucleotides that are associated with at least one GO category. Each nucleotide is counted only once. In “proximity” mode, the background GO frequency computation is similar, but based only on basal and extended regulatory domains.

For each GO category, GOntact will compare the frequency observed in the foreground CRE set with the expected frequency computed based on the background set, using a binomial test. Adjusted p-values that control the false discovery rate are computed with the Benjamini-Hochberg procedure. Output files contain several tab-separated columns, with rows corresponding to GO categories. For each GO category, output files provide its identifier and its description, the number of CREs associated to it in the foreground set, the observed GO frequency in the foreground set, the number of CREs (or nucleotides) associated to it in the background set, the expected GO frequency computed based on the background, the enrichment value (i.e. the ratio between the observed and the expected frequencies), the p-value and the adjusted p-value obtained with the Benjamini-Hochberg procedure.

## Code and data availability

The source OCaml code and the R scripts used to generate graphics and statistical analyses are freely available in a GitLab repository:

https://gitlab.in2p3.fr/anamaria.necsulea/GOntact.

Detailed installation and usage instructions are provided in GitLab. For reproducibility, we provide GOntact binaries in a Docker container. The data used in the manuscript and the results of the GOntact runs analyzed in the manuscript are available here:

http://pbil.univ-lyon1.fr/members/necsulea/GOntact

## Author contribution

A.L., P.V. and A.N. developed the GOntact standalone tool. P.V. developed the GOntact webserver. A.L. and A.N. performed biological analyses. A.L., E.T., P.V. and A.N. designed biological analyses. A.N. wrote the manuscript, with input from all authors.

## Acknowledgements

We thank C. Berthelot, Y. Ghavi-Helm, F. Picard, C. François and D. Mouchiroud for discussions and advice on the project. We thank Stéphane Delmotte for help with webserver deployment. Computational analyses were performed using the computing facilities of the CC LBBE/PRABI and the Core Cluster of the Institut Frangais de Bioinformatique (IFB) (ANR-11-INBS-0013). This work was funded by the French National Research Agency (ANR-17-CE12-0019-01 “LncEvoSys”).

---

**Supplementary Figure 1.**
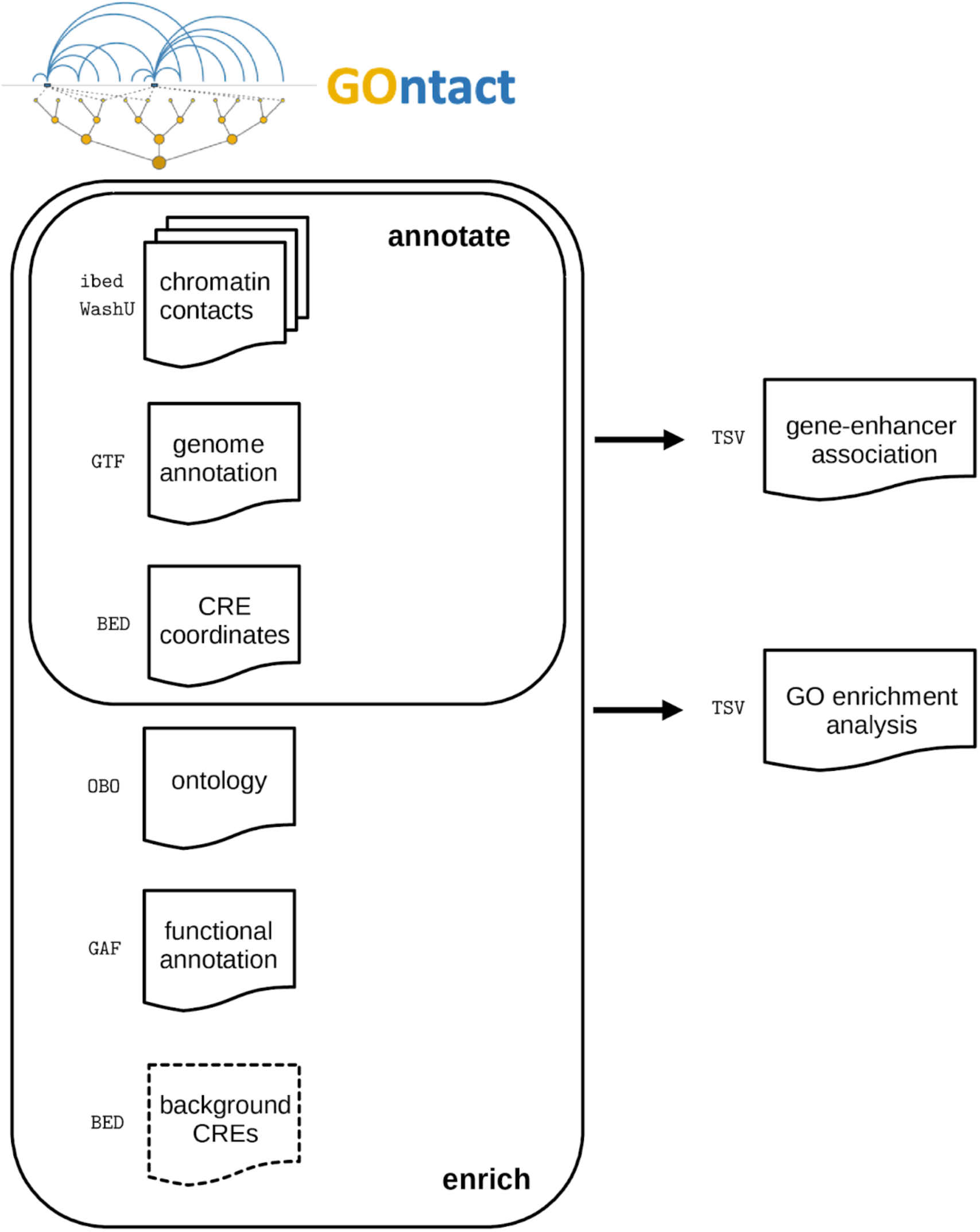
Schematic representation of the input data required by the two GOntact sub-commands (“annotate” and “enrich”) and of their main outputs. Dotted lines depict facultative input files. Chromatin contact data is compulsory only in “contacts” mode. If background sets of CREs are not provided, GOntact uses the whole genome as a background. The two sub-commands (“annotate” and “enrich”) share several parameters and input data types, but can be used independently. The “enrich” subcommand will internally call functions from the “annotate” module to infer gene-CRE target relationships.

**Supplementary Figure 2.**
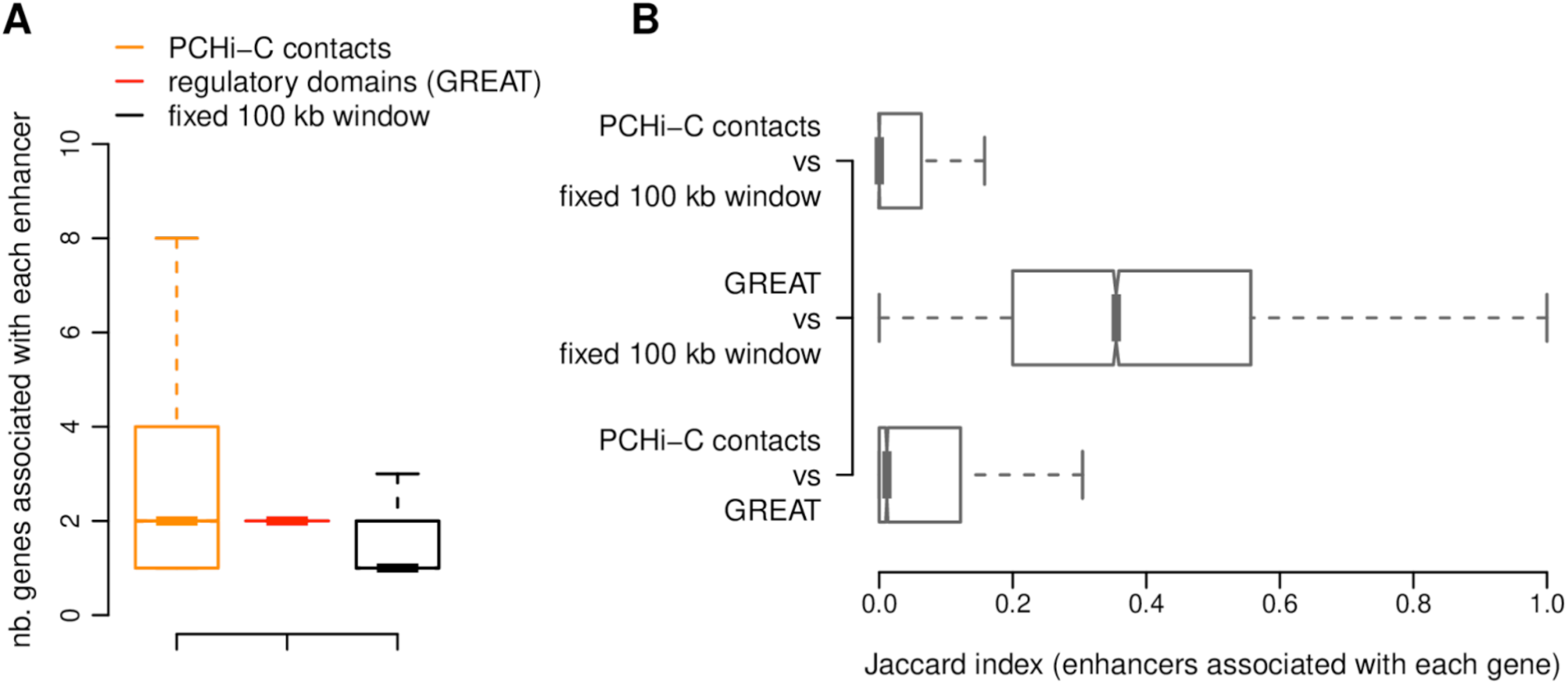
Gene-enhancer associations. **A)** Boxplot representing the number of genes associated with each ENCODE-predicted enhancer. Gene-enhancer associations were predicted based on PCHi-C contacts (orange), regulatory domains (red) or on simple genomic proximity (fixed 100 kb window, black). **B)** Boxplots representing the distribution of the Jaccard indexes obtained through the comparison of sets of enhancers associated with genes for pairs of approaches: PCHi-C contacts versus regulatory domains (bottom), regulatory domains versus fixed 100 kb window (middle), PCHi-C contacts versus fixed 100 kb window (top).

**Supplementary Figure 3.**
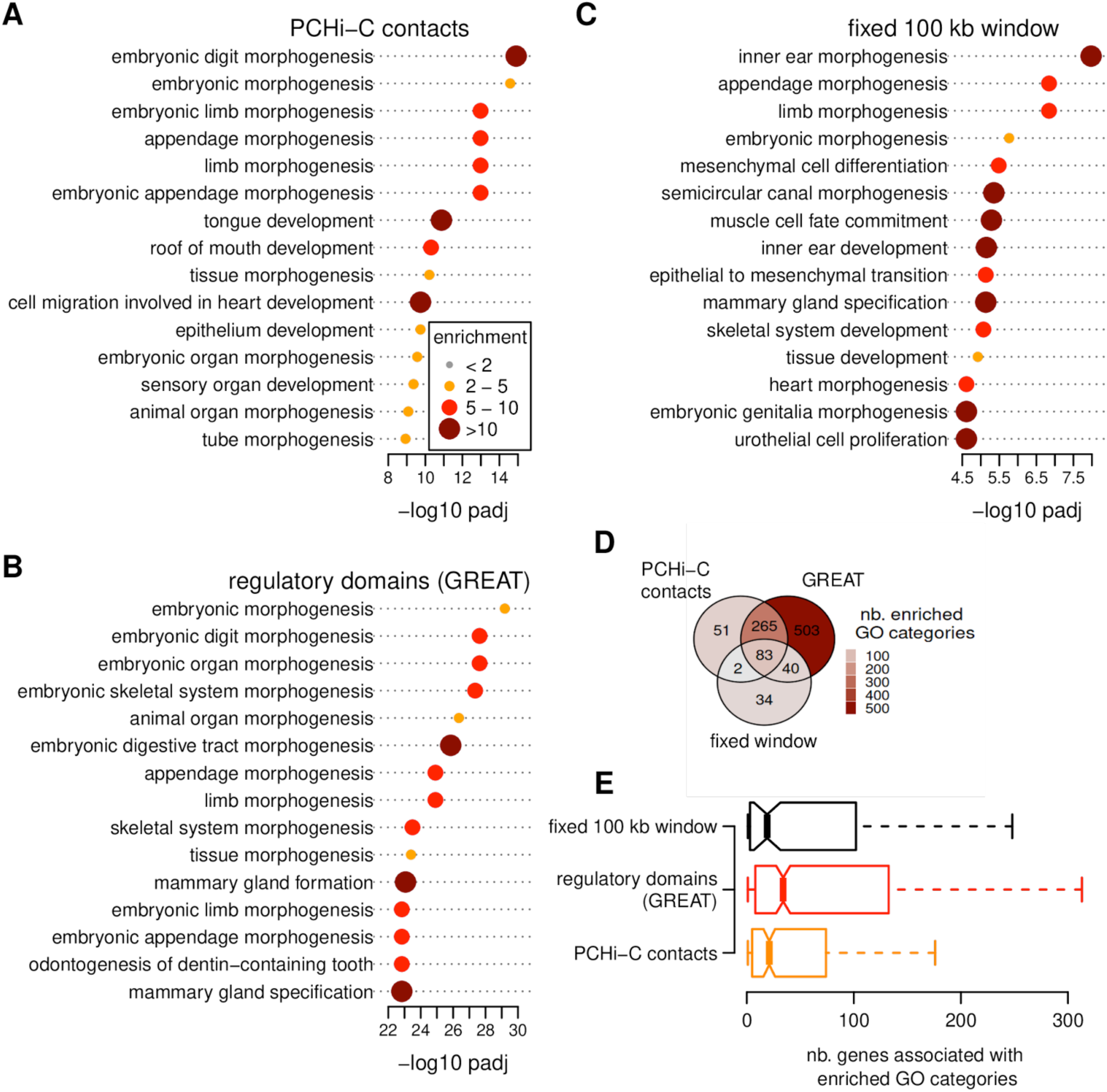
Gene Ontology enrichments for Vista limb enhancers. **A)** Barplot representing log10-transformed adjusted p-values for the top 15 significantly enriched GO categories, for a GO enrichment analysis performed based strictly on PCHi-C chromatin contact (we did not include a basal regulatory domain). Dot sizes and colors represent enrichment values. **B)** Same as **A)**, for the “regulatory domains” approach (McLean et al., 2010). **C)** Same as **A)**, for the simple genomic proximity approach (fixed 100 kb window). **D)** Venn diagram representing the overlap between the GO categories found to be significantly enriched (FDR < 0.01) by the three approaches. **E)** Boxplot representing the number of genes associated with enriched GO categories, for the three methods. For all panels, enrichment analyses are performed using as a background the set of human ENCODE predicted enhancers. Analyses performed using the whole genome as a background are provided in Supplementary Dataset 2.

**Supplementary Figure 4.**
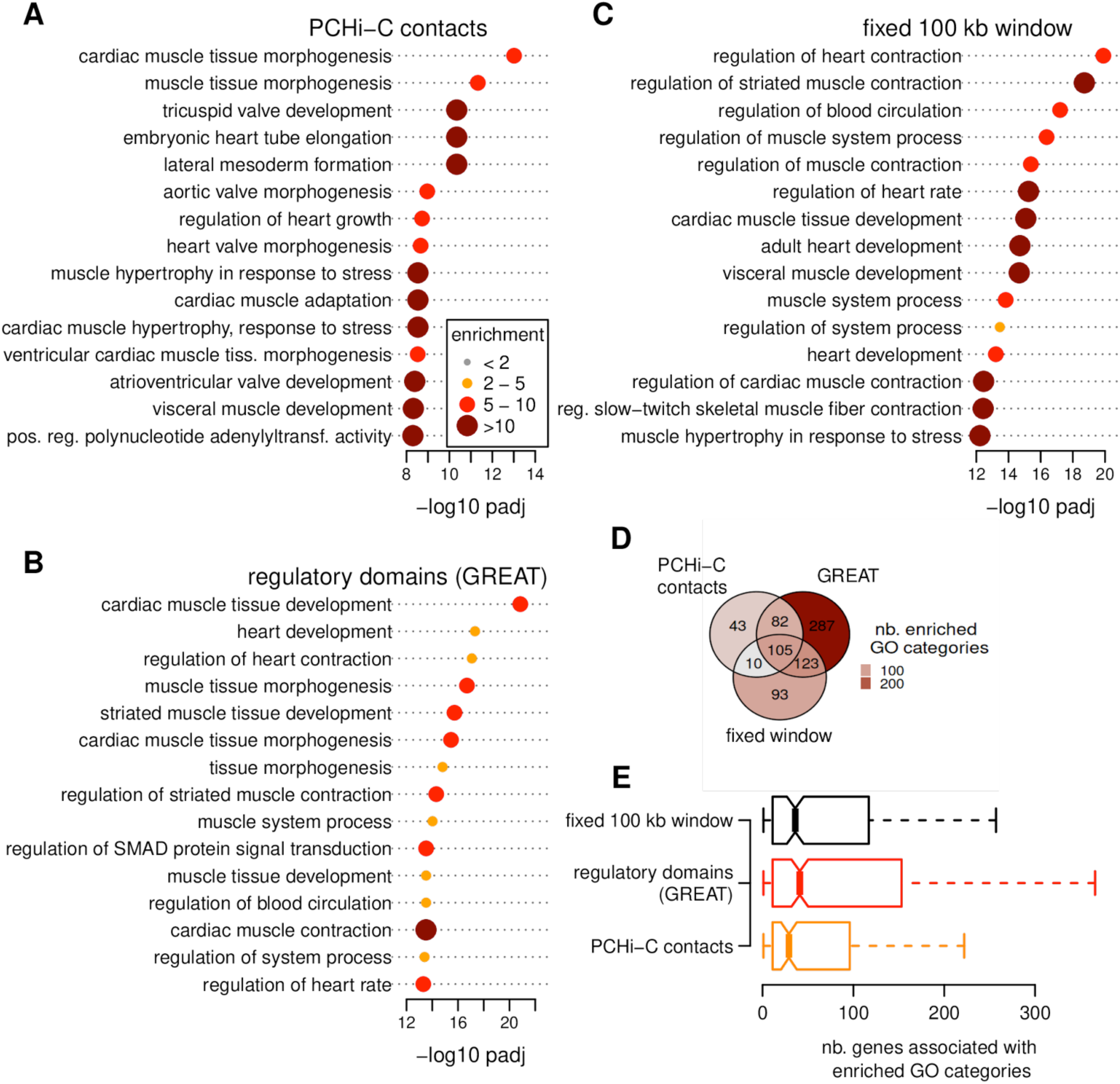
Gene Ontology enrichments for Vista heart enhancers. **A)** Barplot representing log10-transformed adjusted p-values for the top 15 significantly enriched GO categories, for a GO enrichment analysis performed based strictly on chromatin contacts (we did not include a basal regulatory domain). Dot sizes and colors represent enrichment values. **B)** Same as **A)**, for the “regulatory domains” approach (McLean et al., 2010). **C)** Same as **A)**, for the simple genomic proximity approach (fixed 100 kb window). **D)** Venn diagram representing the overlap between the GO categories found to be significantly enriched (FDR < 0.01) by the three approaches. **E)** Boxplot representing the number of genes associated with enriched GO categories, for the three methods. For all panels, enrichment analyses are performed using as a background the set of human ENCODE predicted enhancers. Analyses performed using the whole genome as a background are provided in Supplementary Dataset 2.

**Supplementary Figure 5.**
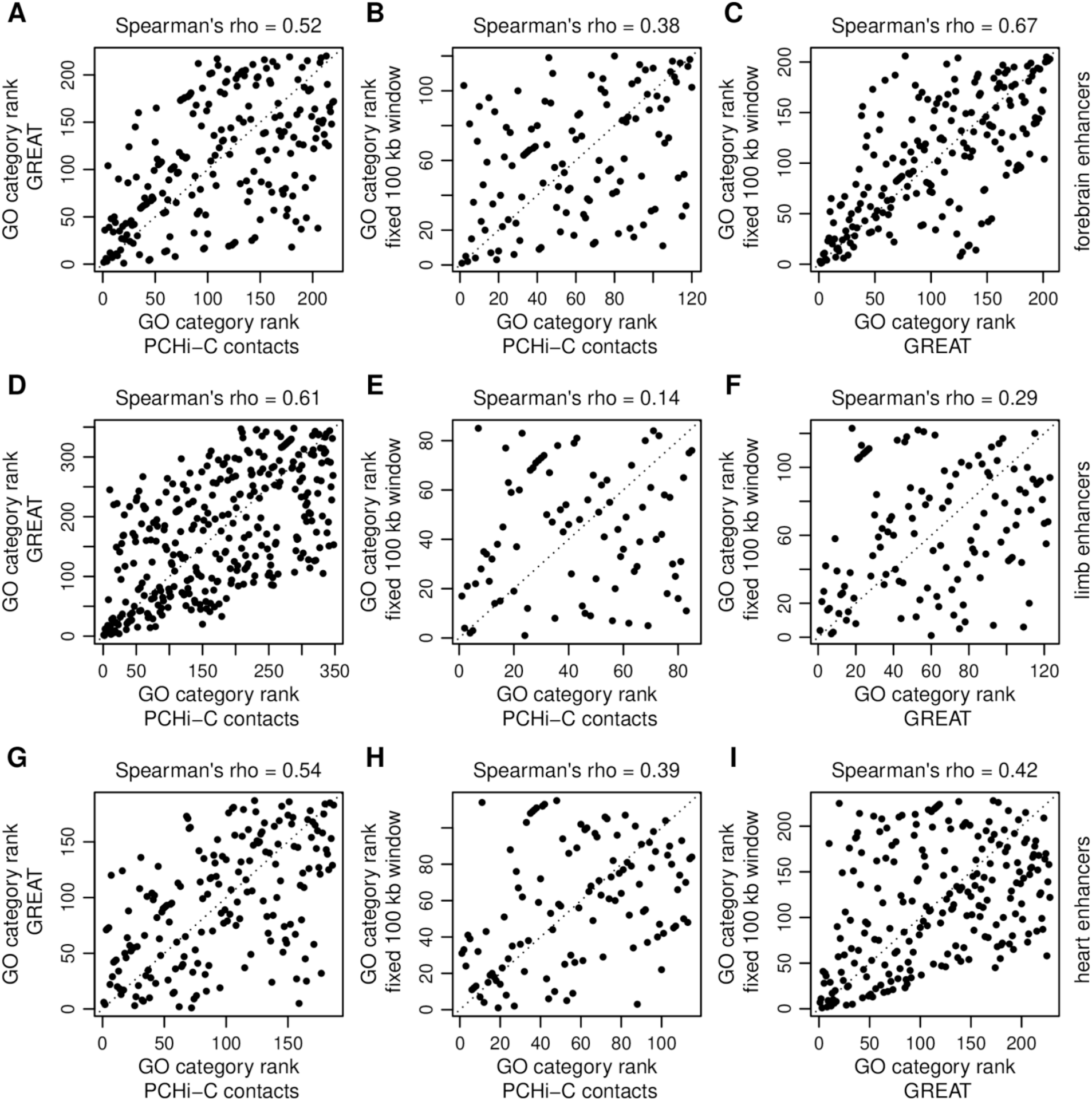
GO categories ranks in enrichment results. **A)** Scatter plot representing a comparison between the ranks of individual GO categories in enrichment analyses (contrasting Vista forebrain enhancers with all ENCODE-predicted enhancers), performed based on PCHi-C contacts (X-axis) and on regulatory domains (Y-axis). Only GO categories found significantly enriched (FDR < 0.01) by both approaches are shown. Each dot is a GO category. **B-C).** Same as **A),** for the two other pairwise comparisons between methods. **D-F)** Same as **A-C)**, for the GO enrichment analysis contrasting Vista limb enhancers with all ENCODE-predicted enhancers. **G-I)** Same as **A-C)**, for the GO enrichment analysis contrasting Vista heart enhancers with all ENCODE-predicted enhancers.

**Supplementary Figure 6.**
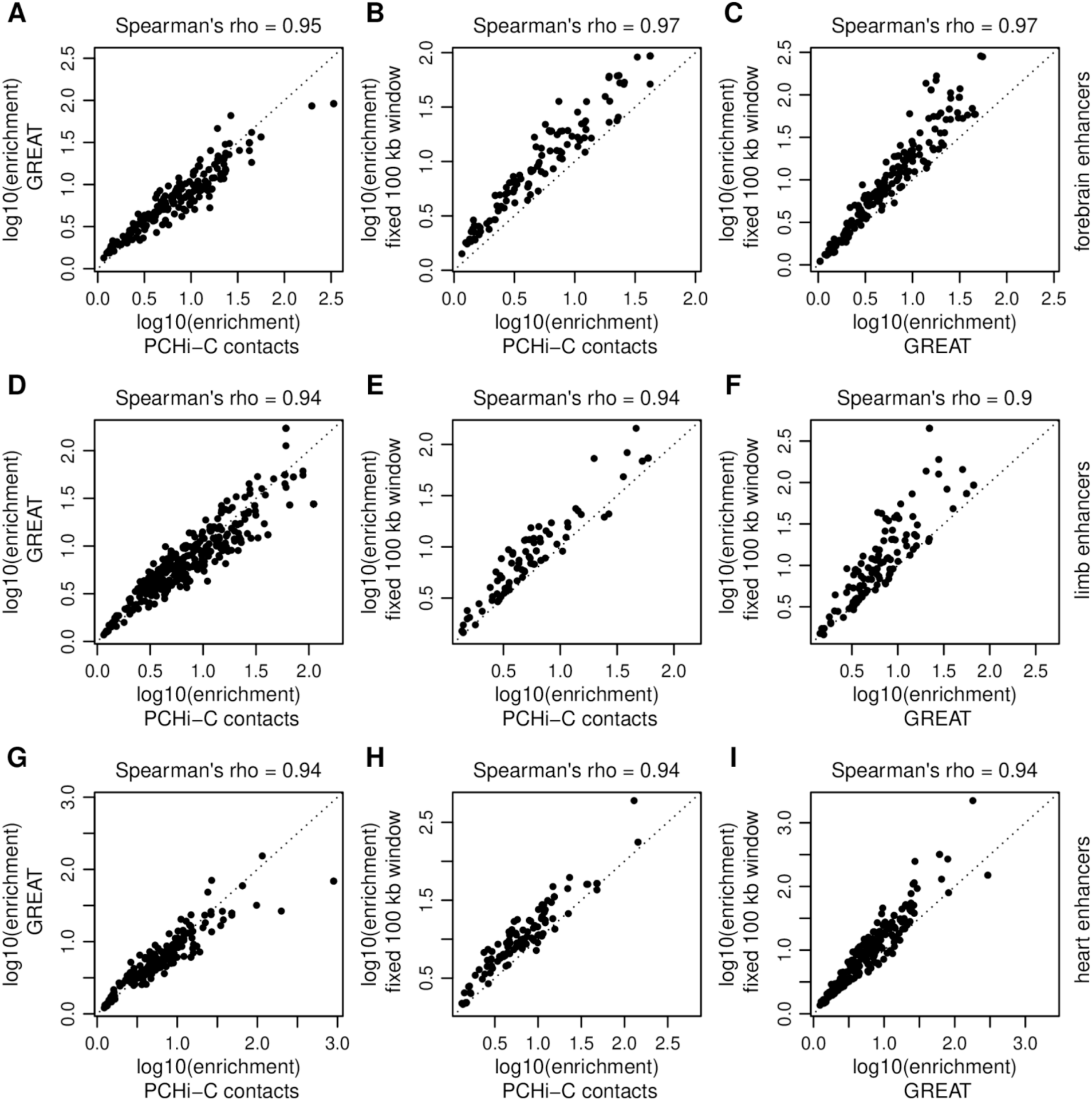
GO categories enrichment values. **A)** Scatter plot representing a comparison between the enrichment values (*i.e.*, the ratio between their frequency in the foreground set and their frequency in the background set) of individual GO categories. This corresponds to enrichment analyses contrasting Vista forebrain enhancers with all ENCODE-predicted enhancers, performed based on PCHi-C contacts (X-axis) and on regulatory domains (Y-axis). Only GO categories found significantly enriched (FDR < 0.01) by both approaches are shown. Each dot is a GO category. **B-C).** Same as **A),** for the two other pairwise comparisons between methods. **D-F)** Same as **A-C)**, for the GO enrichment analysis contrasting Vista limb enhancers with all ENCODE-predicted enhancers. **G-I)** Same as **A-C)**, for the GO enrichment analysis contrasting Vista heart enhancers with all ENCODE-predicted enhancers.

**Supplementary Figure 7.**
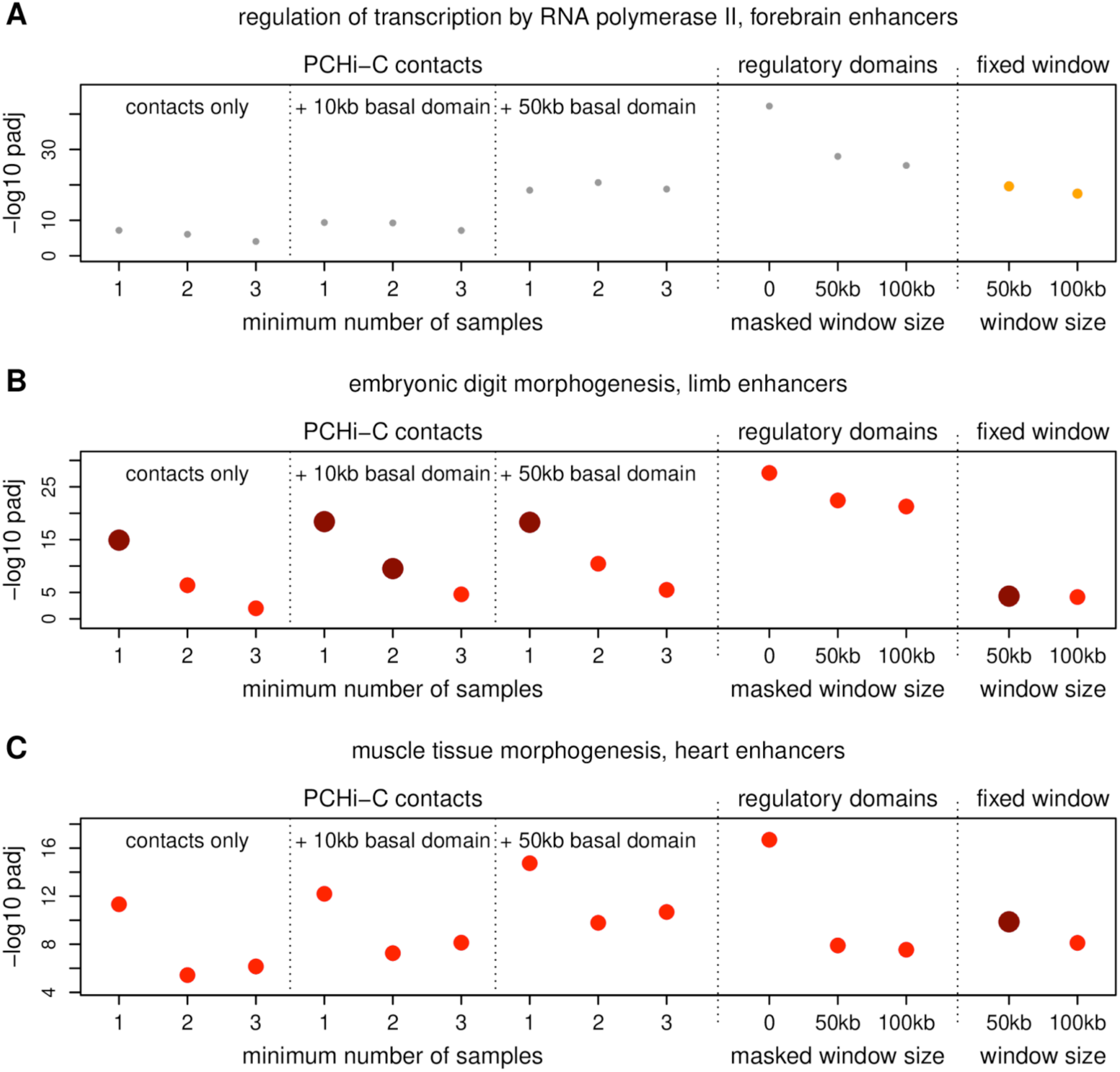
GO enrichment results obtained with different approaches and parameter sets. **A)** Dot plot representing the log10-transformed adjusted p-value for the “regulation of transcription by RNA polymerase II” GO category, for GO enrichment analyses contrasting human forebrain enhancers (Vista) with ENCODE-predicted enhancers. Different columns represent method variants, in order: PCHi-C contacts, requiring that chromatin contacts be observed in at least 1, 2 and 3 samples, respectively; PCHi-C contacts + basal 10 kb domain, with contacts observed in at least 1, 2 and 3 samples; PCHi-C contacts + basal 50 kb domain, with contacts observed in at least 1, 2 and 3 samples; regulatory domains (basal + extended domains, maximum 1 Mb); modified regulatory domains, in which a 50 kb window centered on the TSS is “masked”; modified regulatory domains, in which a 100 kb window centered on the TSS is “masked”; fixed 50 kb window centered on the TSS; fixed 100 kb window centered on the TSS. **B)** Same as **A)**, for the “embryonic digit morphogenesis” GO category, for human limb enhancers. **C)** Same as **A)**, for the “muscle tissue morphogenesis” GO category, for human heart enhancers. Dot sizes and colors represent enrichment values, as in Figure 2.

## Supplementary Material

**Supplementary Dataset 1.** This dataset contains the GOntact runs that were analyzed in this manuscript. This includes the inference of CRE target gene relationships for human ENCODE enhancers (subfolder “CRE_target_genes”) and the GO enrichment analyses for Vista enhancers and hCONDELs (subfolder “GO_enrichment_analyses”), performed with several sets of parameters. The parameters used in the GOntact runs are provided within each subfolder, in a tabulated file named “GOntact_parameters.txt”.

## Notes

### Competing Interest Statement

The authors have declared no competing interest.

### Summary of Updates

This revised version includes substantial updates to the GOntact framework to improve accessibility, speed, and overall usability. The software and interface were optimised to facilitate broader use by the research community. We also expanded the methodological evaluation by benchmarking GOntact on experimental datasets and comparing its results with the existing method GREAT, as well as with alternative approaches and parameter settings. Finally, we added a new application case study analysing Gene Ontology enrichments associated with human-specific conserved element deletions (hCONDELs), illustrating the utility of GOntact for interpreting regulatory genomic variation.

